# Lipid-bound ApoE3 self-assemble into elliptical disc-shaped particles

**DOI:** 10.1101/2020.09.30.319905

**Authors:** Andreas Haahr Larsen, Nicolai Tidemand Johansen, Michael Gajhede, Lise Arleth, Søren Roi Midtgaard

## Abstract

Apolipoproteins are vital to lipid metabolism and cholesterol transport in the human body. Here we present a structural study of the lipid-bound particles formed by ApoE3 in a full-length and a truncated version. The particles are formed with, respectively, POPC and DMPC and investigated by small-angle X-ray scattering and negative stain electron microscopy. We find that lipid-bound ApoE3 particles are elliptical, disc-shaped particles composed of a central lipid bilayer encircled by two amphipathic ApoE3 proteins. We went on to investigate a truncated form of ApoE3 containing only residue 80 to 255 (ApoE3^80-255^), which is the central helical repeat segment of ApoE3. The lipid-bound ApoE3^80-255^ particles are found to have the same morphology as the particles with full-length ApoE3. However, they are larger, and form more heterogeneous discoidal structures with four proteins per particle. This behavior is in contrast to ApoA1 where the highly similar helical repeat domain determines the size and stoichiometry of the formed particles both in the case of full-length and truncated ApoA1. Our data hence points towards different mechanisms for lipid bilayer structural modulation by ApoA1 and ApoE3 due to different roles of the non-repeat segments.

## Introduction

High density lipoprotein (HDL) particles are essential for lipid metabolism and transport in the human body [1], and its protein constituents are named apolipoproteins. Human apolipoprotein E (ApoE) is a 299 amino acid, 34 kDa polymorphic protein that together with ApoA1, ApoA2, ApoA4 are the primary apolipoproteins found in human plasma [2]. In the central nervous system (CNS), ApoE is the main apolipoprotein responsible for transporting cholesterol in HDL-like particles [2,3]. The importance of cholesterol transport in the CNS is highlighted by the fact that ∼20% of all cholesterol in the human body is found in the brain that only accounts for ∼2% of the total body mass. Moreover, ApoE has direct implications in Alzheimer’s disease [4], diabetes [5], and coronary artery disease [6]. ApoE is found in three isoforms, ApoE2, ApoE3 and ApoE4 differing only at residue 112 and 158. ApoE3 is the most common (78% of the general world population) with a cysteine in position 112 and an arginine at position 158 [7]. ApoE4 has arginine at both positions and is less abundant (14%), but is correlated with late-onset Alzheimer disease [8,9] and cardiovascular disease [10]. ApoE2 contains cysteine residues at both positions and is even less common (7%). This isoform correlates with type III hyperlipoproteinemia [6]. Previously, much effort has gone into studying the differences between domains of ApoE3 and ApoE4 [8,11–18]. However, the structure of the full-length protein in association with lipids has been studied less [8,19,20].

ApoA1 and ApoE have very similar amphipathic helical repeat sequences [21,22], and sequence analysis of ApoE reveal eight central 22 amino acid repeats (see www.uniprot.org/uniprot/P02649 under “Family and Domains”). Due to this similarity between ApoA1 and ApoE, we hypothesized that ApoE form discoidal particles resembling the ApoA1-based particles that are commonly referred to as nanodiscs [23–29]. Nanodiscs are composed of a central disc-shaped phospholipid bilayer encircled by a pair of stacked ApoA1 molecules [23,27]. A nanodisc model has previously been proposed for lipid-bound ApoE3 [30], whereas ApoE4-lipid HDL particles have been described as ellipsoid-shaped particles with a twisted bilayer and the ApoE4 organized in a helical hairpin conformation [8,19,20]. Here, we investigate the structure of lipid-bound ApoE3, which is the most common ApoE isoform (78% allele frequency). We test the previously proposed nanodisc model for lipid-bound ApoE3 [30], but use more direct structural data, in the form of small-angle X-ray scattering and electron microscopy, to determine its structure, size and composition.

In lipid-bound ApoA1-based particles, removal of the ApoA1 N-terminal domain does not change the overall morphology of the formed lipid-bound particles [27]. Both full-length and truncated ApoA1 form particles with a well-defined disc morphology. We hypothesized that the same would be true for ApoE3, such that the ApoE3 central amphipathic repeats would be sufficient for disc formation. To investigate whether this was the case, we constructed a truncated version of ApoE3 consisting only of the central eight 22 amino acid helical repeats, residue 80-255 of full-length ApoE3 (ApoE3^80-255^). To our knowledge, this segment has not been studied before.

Here we demonstrate that full-length ApoE3 forms disc-shaped particles when reconstituted with either 1,2-dimyristoyl-*sn*-glycero-3-phosphocholine (DMPC) or with 1-palmitoyl-2-oleoyl-*sn*-glycero-3-phosphocholine (POPC). The DMPC- or POPC-bound ApoE3 particles are structurally similar to lipid-bound ApoA1 particles and self-assemble into disc-shaped particles with an elliptical cross-section. The corresponding DMPC- or POPC-bound ApoE3^80-255^ particles also have a discoidal shape, but they are less homogeneous and larger.

## 1 Materials and methods

### 1.1 ApoE3 production and purification

pET28a(+) plasmids encoding for the ApoE3 and ApoE3^80-255^ sequences (SI-1) coupled to an N-terminal His-tag and a cleavable TEV site were transformed into *E. coli* BL21 star DE3 (Novagen) and plated on LB-agar containing kanamycin. Starter cultures were from a single colony and the next day diluted to an optical density of 600 nm (OD600) = 0.05. Protein expression was induced by the addition of 1 mM IPTG (Fisher) at OD600 = 0.6 and continued for 3h at 37°C. Cells were pelleted by centrifugation, resuspended in buffer A (50 mM Tris-HCl, pH 8, 300 mM NaCl, 20 mM imidazole, 6 M GuHCl), and lysed by freeze/thawing followed by vigorous shaking. Insoluble cell debris was pelleted by centrifugation and the supernatant was loaded on a column containing NiNTA resin (Qiagen) equilibrated in buffer A. The column was washed in three column volumes of buffer A, three column volumes of wash buffer (50 mM Tris-HCl, pH 8, 300 mM NaCl, 40 mM imidazole) including 10 mM sodium cholate, three column volumes of wash buffer, and protein was eluted in buffer B (50 mM Tris-HCl, pH 8, 300 mM NaCl, 250 mM imidazole). For ApoE3^80-255^, 10 mM cholate was included in the second wash and in the elution to suppress aggregation. When appropriate, the N-terminal His-tag was cleaved off by Tobacco etch virus (TEV) protease (TEV S219V mutant produced in-house) added to the protein sample and dialyzed 100-fold against TEV buffer (50 mM Tris-HCl, pH 8, 100 mM NaCl, 1 mM EDTA, 1 mM DTT (+ 10 mM cholate for ApoE3^80-255^)) overnight at ambient temperature. ApoE was separated from the 6xHis-tagged TEV protease and residual His-tagged ApoE on NiNTA resin. Finally, samples were concentrated in vivaspin20 10k molecular weight cutoff (Satorius) concentration filters to 5 mg/mL, flash frozen, and stored at −80 °C until further use.

### 1.2 Self-assembly of the protein/lipid particles

Lipids (either DMPC or POPC) (Avanti Polar Lipids) were mixed with cholate in gel filtration buffer (Tris-HCl pH 7.5, 100 mM NaCl, 1mM DTT), in molar ratio 1:2 (lipid:cholate) and with a final lipid concentration of 50 mM. Protein was mixed with the lipid-cholate solution and diluted with gel filtration buffer to obtain the appropriate protein:lipid ratio and a lipid concentration of 10 mM. Amberlite XAD-2 (Sigma) was added immediately to remove cholate and the samples were incubated overnight at a temperature above the melting transition temperature of the lipid (temperature was 9° C for POPC particles and 28° C for DMPC particles).

### 1.3 Size exclusion Chromatography (SEC)

100-500 µl samples (lipid-bound and lipid-free ApoE3 and ApoE3^80-255^) were applied to an AKTA protein purification system (GE) equipped with a Superdex 200 10/300 (GE healthcare) column and a 500 µl sample loop and run at 0.5 ml/min at 5°C. 250 ul fractions were collected. The investigated ratios were: 70, 90 and 120 POPC per ApoE3; 100 and 150 DMPC per ApoE3; 50, 60 and 100 POPC per ApoE3^80-255^; and 60,70 and 125 DMPC per ApoE3^80-255^.

### 1.4 Sample preparation

Peak fractions from SEC, eluting around 12 ml, were collected and split for analysis by native PAGE, cross-linking, phosphate analysis, EM, and SAXS. The selected ratios were, respectively, 70 POPC and 100 DMPC per ApoE3, and 50 POPC and 60 DMPC per ApoE3^80-255^. For the SAXS measurements and phosphate analysis on POPC-bound ApoE3, the sample with 70 POPC aggregated, so samples with 90 POPC per ApoE3 were used instead. Samples were kept on ice and centrifuged at 14 krpm for 2 min on a mini spin plus centrifuge (Eppendorf) before measurement. The samples for EM were subsequently diluted by addition of gel filtration buffer. Protein concentration was determined by absorption at 280 nm, measured on a Nanodrop 1000 spectrophotometer (Thermo Fisher Scientific).

### 1.5 Native PAGE

Native Polyacrylamide Gel Electrophoresis (PAGE) was performed as previously described [31]. Briefly, samples were mixed with loading buffer and applied to a 4-16% Bis-Tris native gel (Thermo Fisher Scientific) together with NativeMark unstained protein standard (Thermo Fisher Scientific). The gel was subsequently stained with Coomassie blue.

### 1.6 Crosslinking

Samples were buffer exchanged into buffer containing HEPES-NaOH pH 7.5, 100 mM NaCl, 1mM DTT and adjusted to a concentration of 10-20 µM. Freshly prepared BS^3^ (bis-sulfosuccinimidyl-suberate) was added to a final concentration of 0.3 mM and left to incubate on ice for 2 hours. The reaction was quenched by adding cold 1 M Tris-HCl pH 7.5 to the sample to a final concentration of 50 mM. The samples were subjected to SDS-PAGE on 10 % acryl amide gels and stained with Coomassie blue.

### 1.7 Phosphate analysis

The amount of lipids present in the sample was estimated via a colorimetric assay as previously described [32,33]. Briefly, the phospholipids where digested using perchloric acid. The free phosphate was subsequently used to form an inorganic complex with ammonium molybdate that was monitored by UV absorbance at 812 nm. Technical triplicates were performed.

### 1.8 Negative-stain Electron Microscopy (EM)

Copper grids were charged on an Easiglow glow discharger (Ager Scientific) and 2 µl samples with a concentration of ∼0.1 µM, were applied. The samples were subsequently stained three times with uranyl acetate and dried for at least 5 min. EM data for ApoE3 samples were acquired at a Tecnai G2 Spirit TWIN electron microscope (FEI, Thermo Fischer scientific) with varying defocus value of 0.7-1.7 µm at Aarhus University. Images were automatically collected using a Tietz TemCam-F416 CMOS camera at a nominal magnification of 67,000x and a pixel size of 3.14 Å, employing Leginon [34]. EM data for ApoE3^80-255^ samples were collected on a CM100 electron microscope (FEI, Thermo Fischer scientific) with a BioTWIN objective lens and a defocus of 1.7 um at the Core Facility for Integrated Microscopy (CFIM) at University of Copenhagen. Images were collected using an Olympus Veleta camera with magnification 66,000x and a pixel size of 7.7 Å.

### 1.9 Analysis of EM data

Images were converted to *.mcp format and evaluated using XMIPP [35]. The rest of the analysis, from particle picking to 3D model refinement was done in Relion 3.0 [36,37]. Approximately 500 particles were picked manually. The particles were extracted and used for an initial 2D classification, and a selection of the initial 2D classes was used for auto picking of the particles (ApoE3-POPC: 1689 micrographs/96,299 particles; ApoE3-DMPC: 1509/122,019; ApoE3^80-255^-POPC: 403/559,157; ApoE3^80-^255-DMPC: 331/384,994). The picked particles were extracted and grouped into 80 2D classes. A subgroup of these particles was selected for further refinement. 2D classification and selection was done iteratively 3-4 times, where aggregated particles, noise-artefacts, and so-called rouleaux [38] were excluded. The final set of particles (ApoE3-POPC 23,200 particles; ApoE3-DMPC 63,352 particles; ApoE3^80-255^-POPC 348,359 particles; ApoE3^80-255^-DMPC 208,598 particles) was used for further 3D analysis. Top views were overrepresented, so a selection of all side views and a fraction of the top views were used to build an initial model. Using the initial model, 10 3D classes were generated using the whole final set (not only the subset used for the initial model). These classes showed only minor polydispersity. Consequently, all particles were included in the final 3D refinement. The final volume was masked and the resolution was estimated using Fourier Shell Correlation (FSC) and a threshold value of 0.143. The calculated circumference for the ellipse was done using Ramanujan’s approximation.

### 1.10 Small-angle X-ray Scattering (SAXS)

SAXS data were collected at P12, EMBL (Hamburg, Germany), with a wavelength of 1.24 Å and standard settings, giving a q-range of 0.002 Å^-1^ to 0.5 Å^-1^, where *q* = 4*π* sin(*θ*)/*λ*, and 2*θ* is the scattering angle. 20 µl samples were measured at 10 °C using an automatized sample-changer robot [39]. The 2D detector images were azimuthally averaged to yield the 1D scattering curves. 1D curves were buffer subtracted and absolute calibrated using H_2_O to obtain the scattering intensity, *I*(*q*), in units of 1/cm. Data were logarithmically rebinned before further analysis. The pair distance distribution functions, *p(r*), were obtained by Bayesian indirect Fourier transformation (BIFT) [40] as implemented in BayesApp [41], available via GenApp [42] (https://somo.chem.utk.edu/bayesapp/). Data points at *q* > 0.35 Å^-1^ were excluded to avoid fitting the noise in this area. A small constant background was fitted, and the default positivity constraint of the *p*(*r*) was opted out in the BIFT.

### 1.11 SAXS model

The SAXS data were fitted with a model of elliptical nanodiscs, as described in [23,43]. The lipids in the nanodisc model were described as five stacked cylinders, two for the outer lipid head groups, two for the inner hydrophobic tail groups, and one central cylinder for the methyl groups (CH_3_) of the tails. Lipids were surrounded by a hollow cylinder representing the ApoE protein. The molecular volumes (*V*), scattering lengths (*b*), scattering length densities (*ρ*=*b*/*V*) and excess scattering length densities (Δ*ρ=ρ*-*ρ*_*H20*_) of the lipid head and tail groups, methyl groups, proteins and the aqueous buffer are given in Table 1. The volume of the lipids was refined with a few percent in the fitting procedure, which effectively lead to small changes of the resulting *ρ* and Δ*ρ* values (Table 5).

**Table 1.** Input values for SAXS analysis. Molecular volumes (*V*), scattering lengths (*b*), scattering length densities (*ρ*), and excess scattering length densities (Δ*ρ*) for each component of the nanodiscs and for the H_2_O-based solvent. *Lipid volumes were refined within a few percent in the fitting process (Table 5).

| Component | $V$ [Å<sup>3</sup>] | $b$ [10<sup>-11</sup> cm] | $\rho$ [fm/Å<sup>3</sup>] | $\Delta\rho$ [fm/Å<sup>3</sup>] |
| --- | --- | --- | --- | --- |
| PC (lipid head)* | 319 | 4.62 | 1.45 | 0.51 |
| PO (lipid tail)* | 818.8 | 6.71 | 0.82 | -0.12 |
| DM (lipid tail)* | 682.4 | 5.41 | 0.79 | -0.15 |
| CH<sub>3</sub> (lipid tip)* | 108.6 | 0.51 | 0.47 | -0.47 |
| POPC (total)* | 1246.4 | 11.8 | 0.95 | 0.01 |
| DMPC (total)* | 1110.0 | 10.5 | 0.95 | 0.01 |
| ApoE3 | 42291 | 517.8 | 1.22 | 0.28 |
| ApoE3<sup>80-255</sup> | 24494 | 301 | 1.23 | 0.29 |
| H<sub>2</sub>O | 30.0 | 0.28 | 0.94 | 0.0 |

**Table 2.** Results of the phosphate analysis. Standard deviations from technical triplet replicates are given. Three samples were made from each protein prep for the assay. Lipids per particle assume two proteins per particle in ApoE3-POPC and ApoE3-DMPC, and four proteins per particle in ApoE3^80-255^-POPC, according to the nanodisc models that best fitted SAXS data (see later).

| Sample | Lipid per protein | Lipids per particle |
| --- | --- | --- |
| ApoE3-POPC | 90 ± 5 | 180 ± 10 (2 ApoE3/particle) |
| ApoE3-DMPC | 111 ± 2 | 222 ± 4 (2 ApoE3/particle) |
| ApoE3<sup>80-255</sup>-POPC | 80 ± 7 | 320 ± 28 (4 ApoE3<sup>80-255</sup>/particle) |
| ApoE3<sup>80-255</sup>-DMPC | 74 ± 4 | 296 ± 16 (4 ApoE3<sup>80-255</sup>/particle) |

**Table 3.**
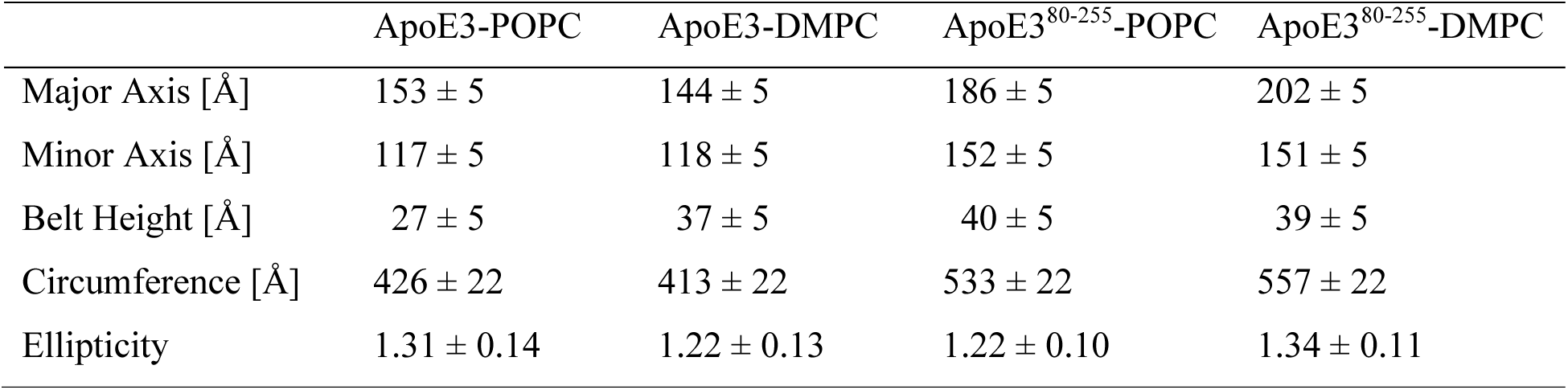
Dimensions from the EM 3D volumes. As measured from the 3D density map, with an estimated uncertainty of ± 5 Å.

**Table 4.** Parameters resulting from indirect Fourier transformation of the SAXS data. Errors are obtained by Bayesian interference [40,41].

| | $D_{\max}$ [Å] | $R_g$ [Å] |
| --- | --- | --- |
| ApoE3-POPC | $181 \pm 6$ | $63.7 \pm 0.3$ |
| ApoE3-DMPC | $151 \pm 6$ | $51.9 \pm 0.2$ |
| ApoE3<sup>80-255</sup>-POPC | $195 \pm 5$ | $64.4 \pm 0.3$ |
| ApoE3<sup>80-255</sup>-DMPC | $192 \pm 7$ | $70.6 \pm 0.5$ |

**Table 5.**
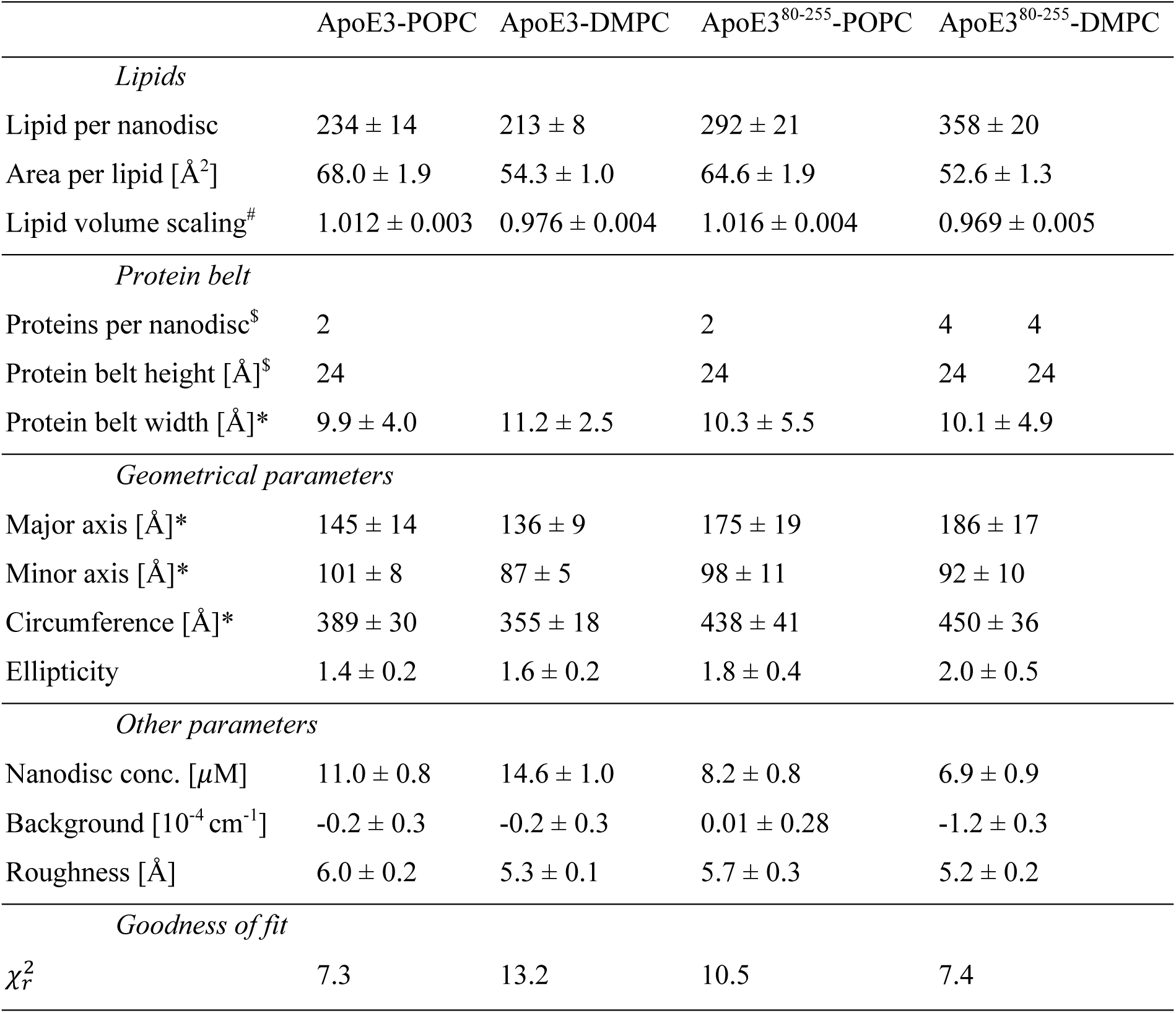
Fitted and derived parameters from the modelling of the SAXS data. Errors on the parameters were retrieved from the covariance matrix and represent one standard deviation. Errors on the derived parameters were calculated by error propagation. ^#^This factor was multiplied with the lipid volumes in Table 1 to obtain the refined lipid volumes. ^$^Fixed model parameter. ^*^Derived from the model parameters.

The goodness of fit of the SAXS model was assessed by the reduced χ^2^. The degrees of freedom were determined from the number of data points (*N*) and the number of fitted parameters (*K*) as *f* = *N*-*K*:

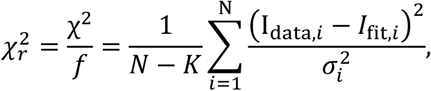

where *I*_*data*_is the experimental data, *σ* is the standard deviation, and *I*_*fit*_ is the fitted intensity. There were seven refined parameters in the model (*K* = 7): number of lipids per nanodisc, area per lipid, scaling of lipid volume, ellipticity of nanodisc, protein concentration, constant background, and roughness [43] (Table 5).

## 2 Results

### 2.1 Self-assembly of lipid bound ApoE3 and ApoE3^80-255^

In order to investigate the response of the two types ApoE proteins to lipids, protein was reconstituted with different amounts of either POPC or DMPC, self-assembled by the cholate assisted method [44] and applied to a gel filtration column (Figure 1). Relevant samples were also subjected to native PAGE or crosslinked and analyzed by SDS-PAGE (Figure 2).

**Figure 1.**
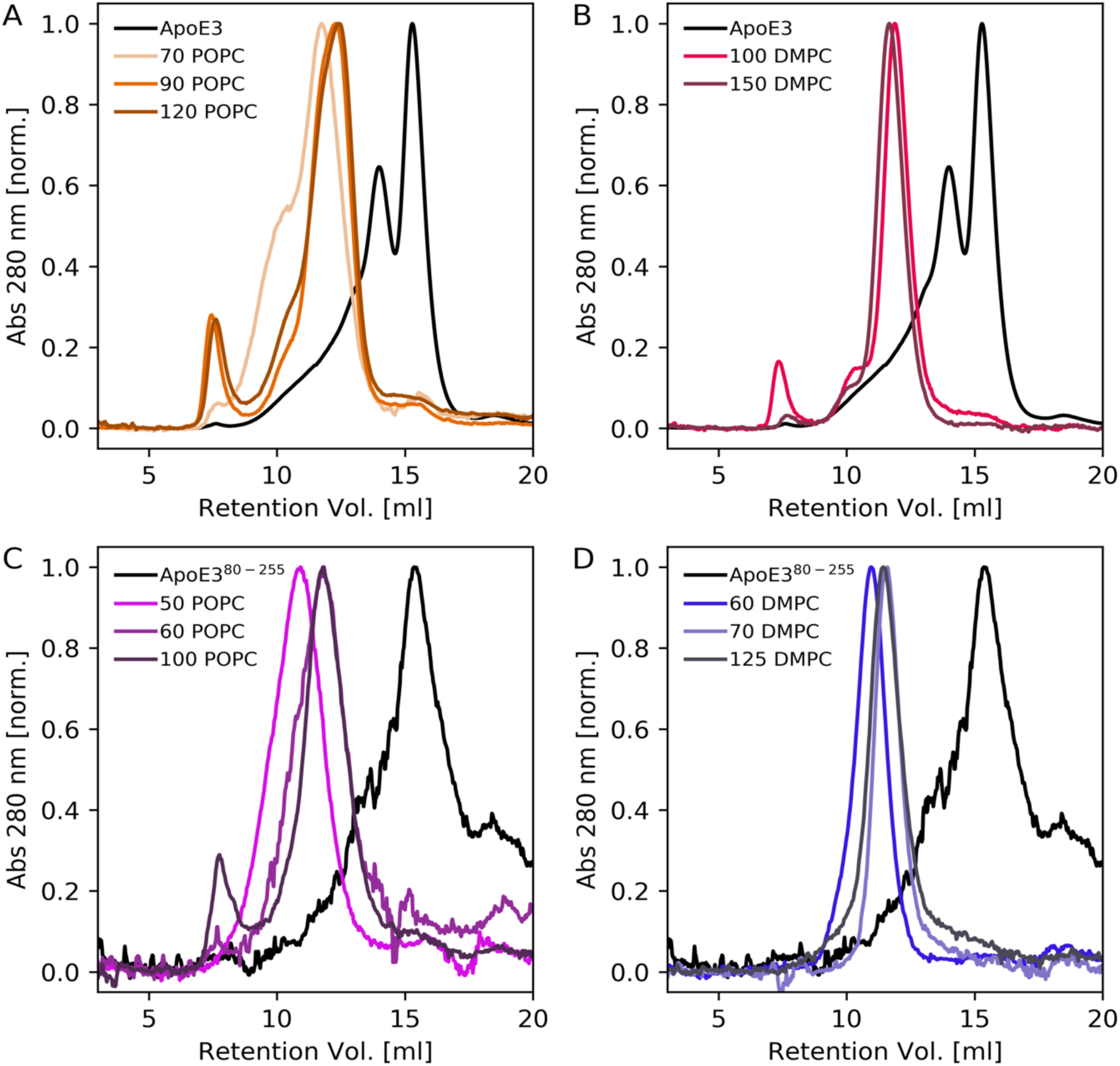
SEC chromatograms with different amounts of lipids per protein. Absorbance is normalized to a peak value of unity. Black lines indicate the pure protein while the numbers of the colored lines indicate the lipid:protein stoichiometry used in the reconstitution (A) ApoE3-POPC, (B) ApoE3-DMPC, (C) ApoE3^80-255^-POPC, and (D) ApoE3^80-255^-DMPC.

**Figure 2.**
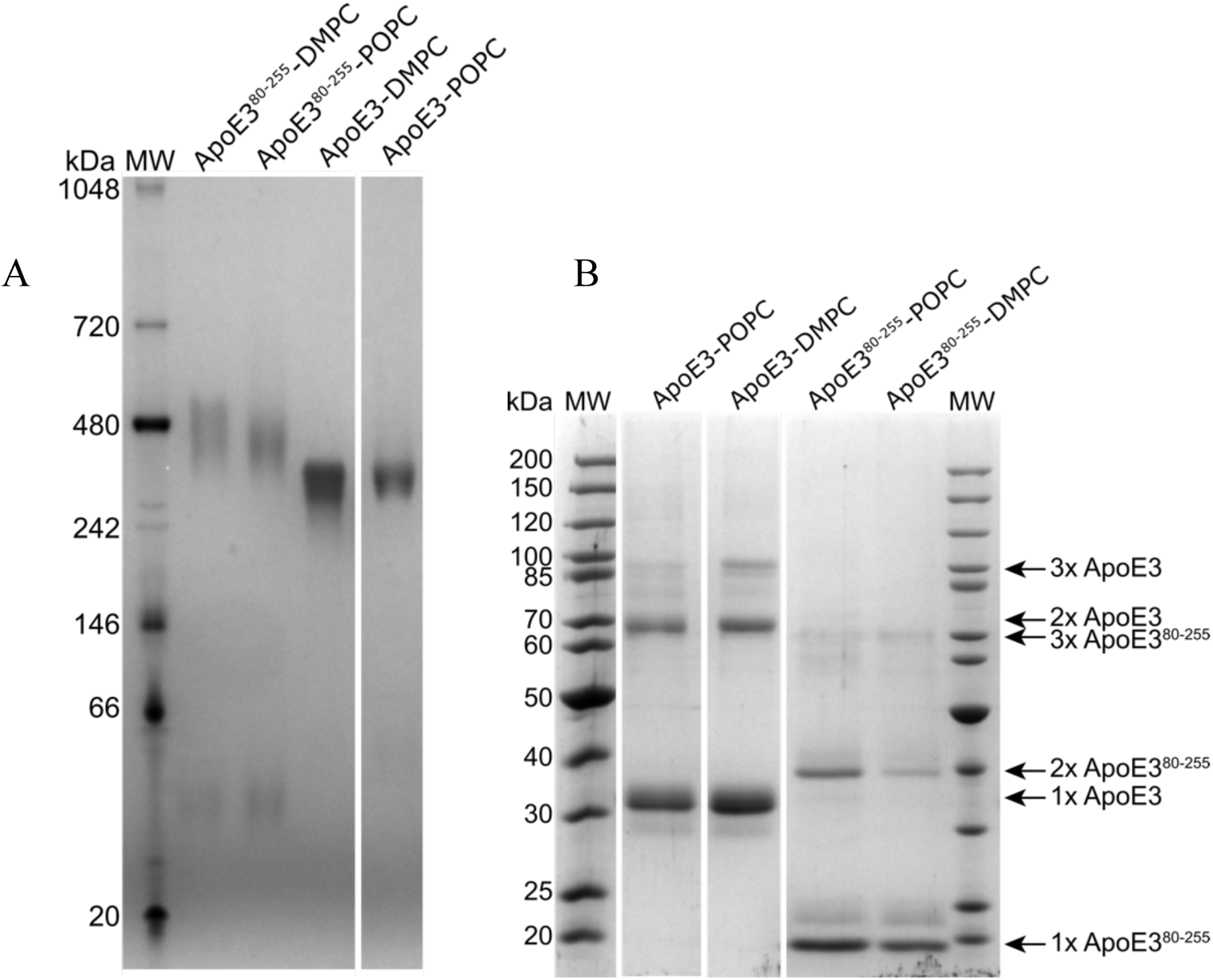
Chemical analysis. The loaded samples are identical to those analyzed by SAXS and phosphate analysis. (A) Blue-Native PAGE analysis of DMPC and POPC-bound ApoE3 and ApoE3^80-255^ particles. (B) Chemical cross-linking of DMPC and POPC-bound ApoE3 and ApoE3^80-255^ particles with Bissulfosuccinimidyl suberate. Cross-linked samples were analyzed by SDS-PAGE. Arrows indicate the protein multimerization number estimated from the molecular weight ladder (MW). All lanes originate from the same gel.

ApoE3 without lipids forms multiple species in solution (Figure 1). This is consistent with previous reports of concentration dependent oligomerization of ApoE3 without lipids [45]. The main peak of ApoE3^80-255^ (176 amino acids) without lipids elutes at approximately the same volume (15.2 ml) as ApoE3 (299 amino acids), hinting to a potential dimerization of this much smaller protein.

The size of all protein-lipid particles depends on the lipid:protein stoichiometry (Figure 1), but surprisingly, the ApoE3^80-255^-lipid particles are larger than the ApoE3-lipid particles (Figure 1 and Figure 2A). ApoE3-based particles have more narrow SEC peaks than ApoE3^80-255^ -based particles. This is also reflected in more narrow bands in the native PAGE analysis (Figure 2A). We note that the SEC distributions are slightly narrower when the particles are produced with DMPC than with POPC (Figure 1). We also note that SEC was conducted at 5 °C where DMPC is in the gel phase, and POPC is in the fluid phase.

### 2.2 Protein:lipid stoichiometry in the self-assembled particles

Phosphate analysis of ApoE3-lipid particles yielded, respectively, 90 POPC or 111 DMPC molecules per protein (Table 2), whereas ApoE3^80-255^-based particles had 80 POPC or 74 DMPC molecules per protein (Table 2). I.e. there were most lipids per protein in the ApoE3-based particles. Since these were also smaller, as seen from SEC and native PAGE, we hypothesized that there were more proteins per particle in the ApoE3^80-255^-based HDL particles than in the ApoE3-based particles. This was further investigated by chemical crosslinking of the proteins (Figure 2B).

Crosslinking experiments (Figure 2B) revealed a large population of monomeric and dimeric ApoE3, both with POPC and DMPC. A weaker band with trimeric ApoE3 was also observed. Crosslinking for lipid-bound ApoE3^80-255^ likewise results in clear bands for monomeric and dimeric ApoE3^80-255^ and a weak band for trimeric ApoE3^80-255^. The monomeric band is probably due to incomplete cross-linking and is hence consistent with particles with 1, 2, or 3 ApoE3. The number of lipids per protein from the phosphate analysis (Table 2), held up against the relative sizes from native PAGE and SEC points towards a larger protein:lipid stoichiometry for ApoE^80-255^ than for ApoE3. This is not observed in the crosslinking analysis. To investigate the overall particle morphology with minimum assumptions, we imaged the particles with EM.

### 2.3 Overall discoidal particle structure revealed by EM

On the 2D EM images, it is clear that the HDL particles exhibit an overall discoidal morphology, for both full-length and truncated ApoE3 with POPC or DMPC (Figure 3). Most images of single particles were top views (Figure 3 and 4), as a majority of the side views are stacks or so-called rouleaux of discoidal particles (Figure 3). Rouleaux have also been observed by others [8,46], and have been attributed to be artifacts from the negative staining and immobilization on a grid surface [8]. Unfortunately, the rouleaux could not be used further in the single-particle structure refinement. The single particles, excluding rouleaux, were refined to 2D classes, and the variety within these 2D classes show that the particles have some ellipticity with a both size and shape polydispersity (Figure 4).

**Figure 3.**
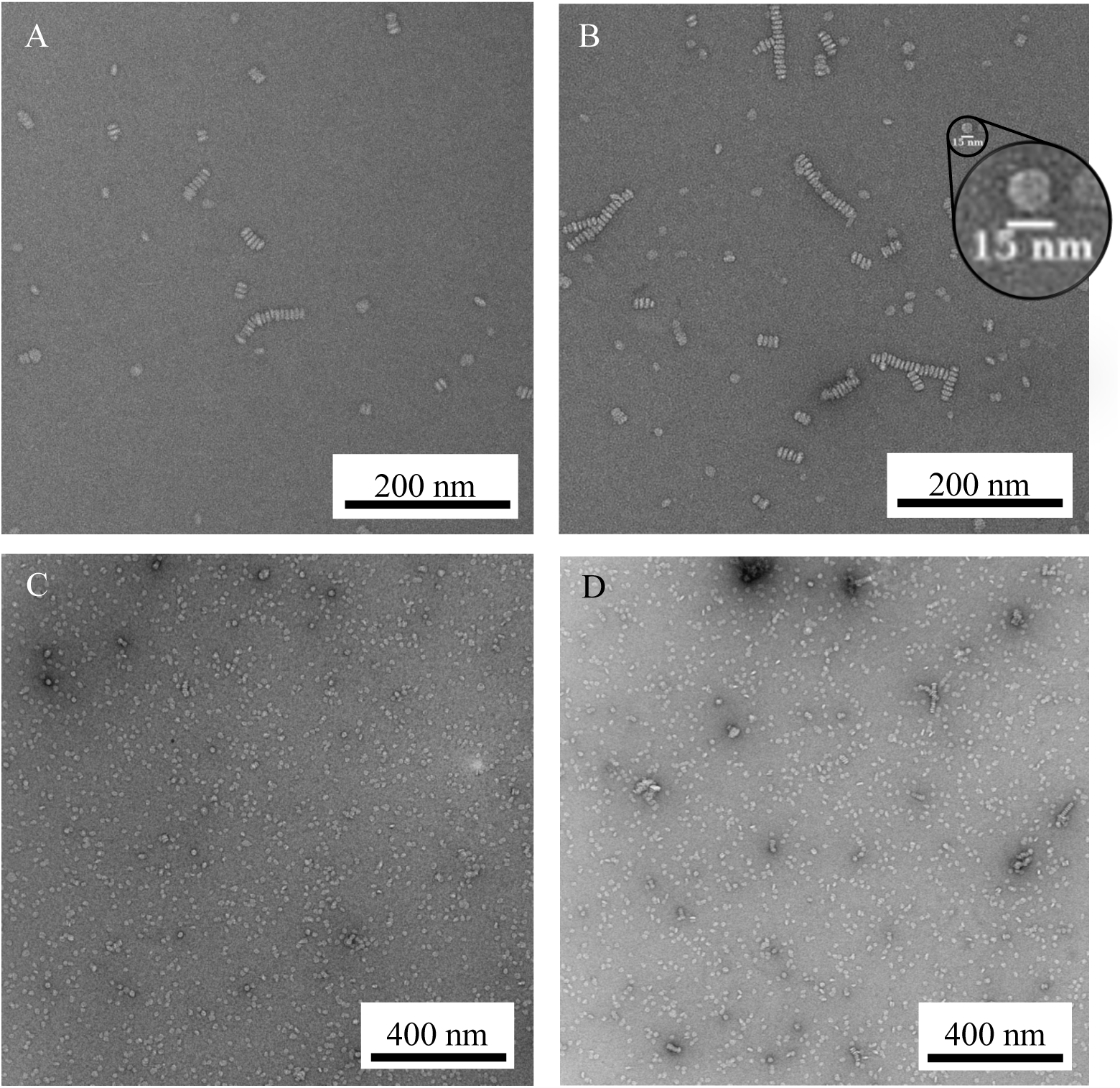
Representative negative stain EM micrographs. (A) ApoE3-POPC, (B) ApoE3-DMPC, with inserted additional 15 nm scale bar for one representative particle, (C) ApoE3^80-255^-POPC, and (D) ApoE3^80-255^-DMPC.

**Figure 4.**
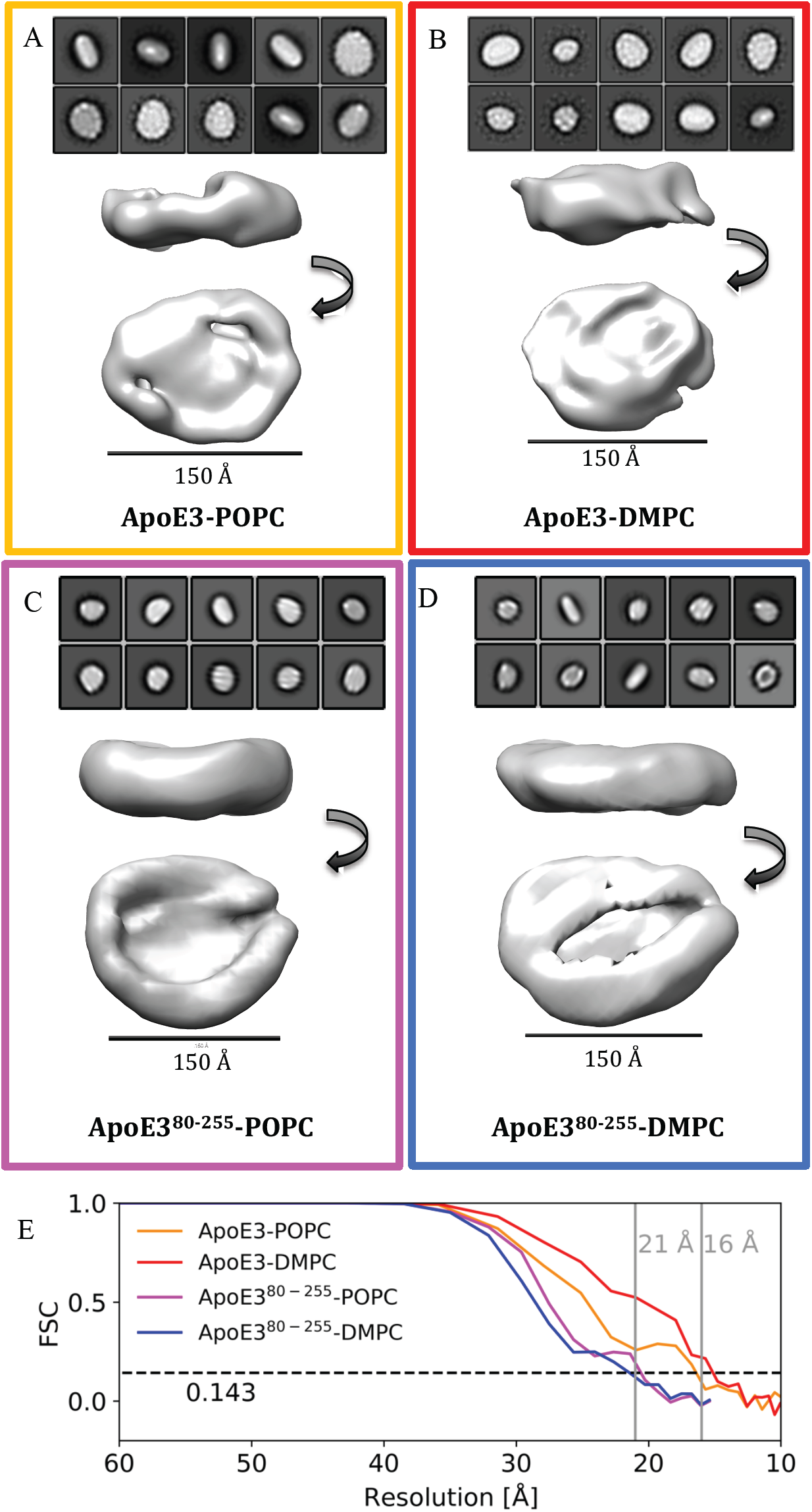
Classes from negative stain EM. (A-D) The ten most populated 2D classes shown together with the resulting 3D reconstruction. 2D classes for ApoE3-lipid and ApoE3^80-255^-lipid particles are not on the same scale as they were measured on different instruments with different settings. (E) Fourier shell correlation curves for all 3D classes.

3D classification was performed from the 2D classes, and for each sample, a volume was determined (Figure 4). ApoE3-lipid particle volumes had major and minor axes of about 150 Å and 120 Å, and a resulting ellipticity of about 1.2 to 1.3 (Table 3). The particles are slightly larger than previously reported [8,20]. The resolution of the refined 3D density maps for ApoE3 was about 16 Å (Figure 4). The ApoE3^80-255^ volumes were larger, with major and minor axes of about 200 Å and 150 Å and an ellipticity between 1.2 and 1.3 (Table 3) with a resolution estimated to be 21 Å (Figure 4). The lipid core of the 3D structures generally had lower density, whereas the protein rim was more clearly resolved (Figure 4). This may be explained by uranyl binding to the lipid bilayer [47,48], which decreases the contrast of the lipid core.

The EM data thus reveal that the HDL particles with ApoE3 or ApoE3^80-255^ are disc-shaped. The particles with ApoE3^80-255^ are larger than particles with ApoE3. This is consistent with the combined native page and phosphate analysis. Again, this indicates that there are more protein per lipid in the particles with ApoE3^80-255^, despite the crosslinking results. We used SAXS to further investigate the shape and stoichiometry of the formed particles under solution conditions.

### 2.4 Visual inspection and model-free analysis of the SAXS data

To examine the structure in more detail, we measured all samples using SAXS (Figure 5). This method has the advantage of probing the solution structure of the particles without exposing them to a heavy atom salt and without fixation to a grid. In addition, SAXS can provide more detailed information about the stoichiometry and shape of the particle.

**Figure 5.**
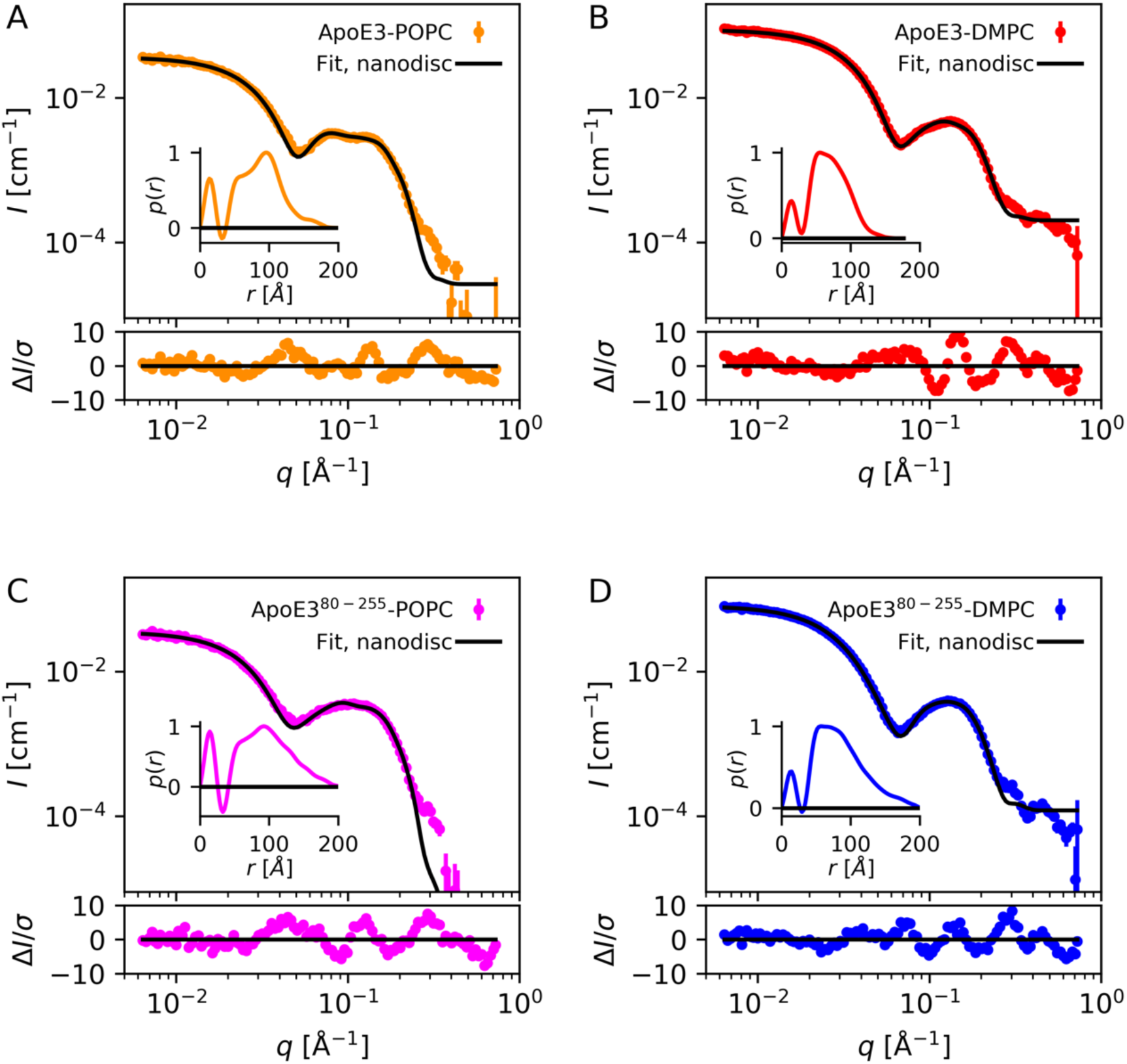
SAXS data and model fits. (A) ApoE3-POPC. (B) ApoE3-DMPC. (C) ApoE3^80-255^-POPC. (D) ApoE3^80-255^-DMPC. Residual plots included to evaluate fit quality. Inserts are the resulting *p*(*r*) functions.

A Guinier plot of the SAXS data exhibit a linear slope at low *q* values (Figure SI-1), showing that the samples are not aggregated.

The characteristic minima in the SAXS curves around 0.05 Å^-1^ for POPC and 0.07 Å^-1^ for DMPC are also observed in scattering patterns from ApoA1-lipid particles [27], ApoE3-lipid particles [20], and MSP1D1-based nanodisc particles [23] and is indicative of particles with a core-shell contrast. For these particles, the core-shell contrast stems from the negative excess scattering length density (Δ*ρ*) of the lipid hydrophobic tails (Table 1) while both the protein belt and the lipid head groups have positive Δ*ρ*. Furthermore, the wide peak around 0.1 Å^-1^ has a double bump when POPC is used (Fig 6A), which is similar to the scattering patterns from related ApoA1-POPC discoidal particles [27]. The pair distance distribution functions, *p(r)*, have a negative contribution at around 30 Å (Figure 5, inset). This is a result of the oscillating contrast and is again characteristic for SAXS data from lipid bilayer-containing particles. The ApoE3-DMPC particles show an overall maximum size, *D*_*max*_, of ∼150 Å and an *R*_*g*_ of ∼52 Å (Table 4). This is in agreement with previously published results on lipid-bound ApoE4 [20]. The ApoE3-POPC particles have a larger maximum size of ∼180 Å and an *R*_*g*_ of ∼64 Å (Table 3). As *R*_*g*_ depends on the contrast and hence the type of lipid used, it is not possible to directly compare this value with the previous values obtained for lipid-bound ApoE4 with 1,2-dipalmitoyl-sn-glycero-3-phosphocholine (DPPC) [20]. Lipid-bound ApoE3^80-255^ particles have larger *D*_max_ of almost 200 Å and *R*_*g*_ of 60-70 Å. So, even before modelling, the SAXS data reveal that ApoE3^80-255^-based particles are significantly larger than ApoE3-based particles, which is consistent with EM, native PAGE, phosphate analysis and SEC. We further used model-based analysis to obtain detailed information about the shape and stoichiometry of the discoidal particles.

**Figure 6.**
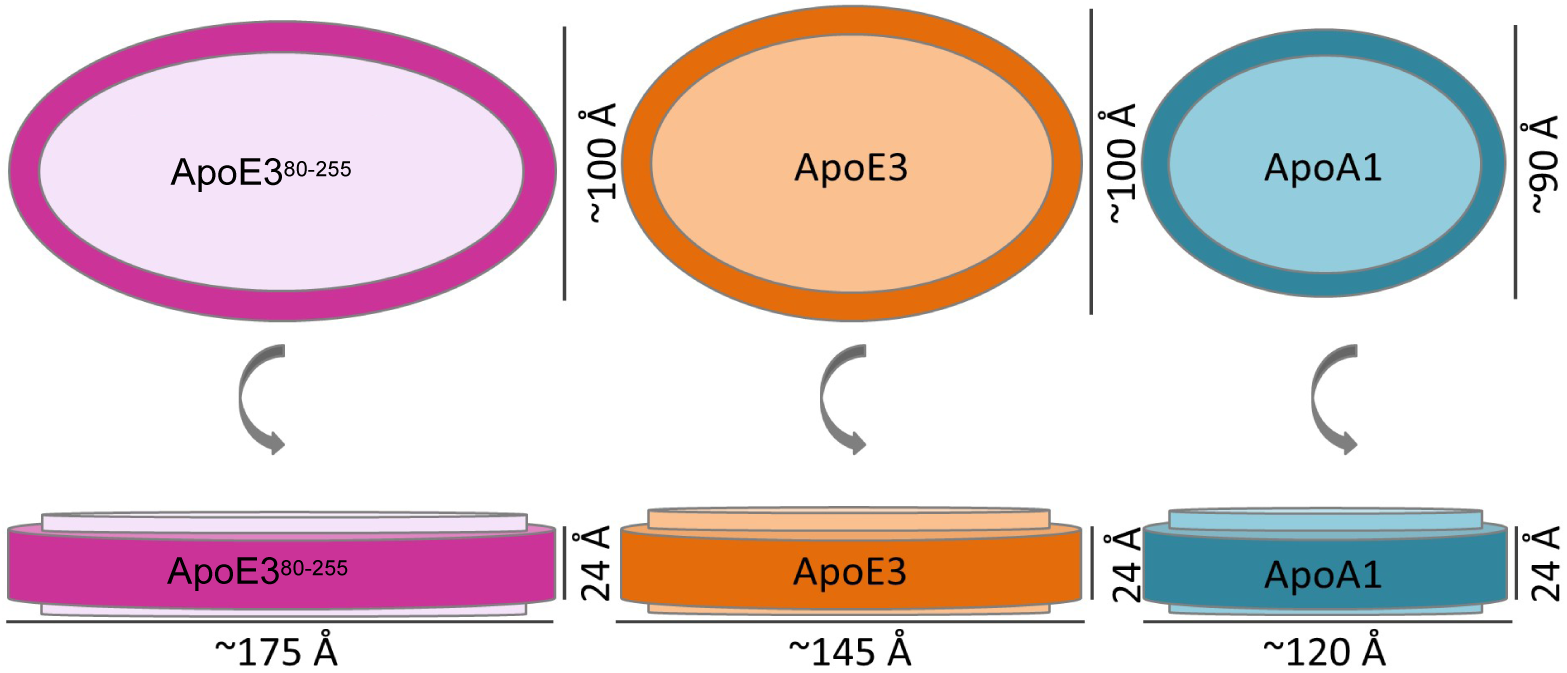
Schematic illustration of the refined structures. Dimensions from the SAXS analysis. ApoA1-POPC dimensions from [27] shown for easy comparison.

### 2.5 Discoidal model is consistent with SAXS data

A discoidal model similar to the EM envelope was fitted to the SAXS data. The model is based on the one we have previously used to describe ApoA1-based nanodiscs [23,27,43] and as in this previous work, molecular constraints (molecular volumes and scattering lengths of the protein belts and the lipid head and tail groups) were included to keep the model consistent with prior knowledge and avoid unphysical solutions. The model described an elliptical lipid bilayer disc surrounded by a protein belt (Figure 6). The refined fitting parameters are given in Table 5.

For lipid-bound ApoE3 particles, we obtained the best model fits when applying a model with two proteins per particle. The number of lipids per nanodisc were refined to 234 ± 14 POPC or 213 ± 8 DMPC lipids, which is consistent with the phosphate analysis (Table 2), when taking into account the simplicity of the SAXS model and uncertainty of both experimental techniques. The two-protein nanodisc model described the data well in the full *q*-range (Figure 5) and the fitted and derived parameters were refined to physically reasonable values (Table 5).

The model was not perfect, as revealed by the residuals: for a perfect model fitted to ideal data, 99.7 % of the residuals would lie in between −3 and 3, and we observed residuals with nominal values up 10. This discrepancy is likely due to the minor trimeric population revealed by cross-linking, as well as a more general heterogeneity also observed in EM. It is, however, striking that a simple geometric model with strict molecular constraints captures the experimental data so well.

The SAXS data of ApoE3^80-255^ with POPC or DMPC were fitted very poorly by the nanodisc model with two protein per disc (fit not shown). Thus, we excluded this model. The data were, however, fitted much better using a model with four proteins per disc (Figure 5). The 4-protein nanodisc models contained respectively 292 ± 21 POPC or 358 ± 20 DMPC lipids per nanodisc, which are consistent with the phosphate analysis (Table 2).

The area per lipid headgroup is a good check of whether the applied model and refined values are realistic. The area per lipid were found to be 65-68 Å^2^ for POPC and 53-54 Å^2^ for DMPC (Table 5). POPC was in the fluid pahse in the SAXS experiment, whereas DMPC was in the gel phase. Therefore, the observation are in line with our expectation. These areas also agree well with previously reported values for other types of nanodiscs with POPC [23,27,49] and DMPC [50,51].

## 3 Discussion

### 3.1 Lipid-bound ApoE3 form heterogeneous discoidal particles with two proteins per particle

EM data showed that ApoE3 form discoidal particles. SAXS data and phosphate analysis supported a nanodisc model containing 2 proteins and ∼200 DMPC or POPC lipids. Crosslinking suggested that there was an additional minor population of particles with 3 proteins per disc, and together with the heterogeneity, which was observed in EM, this gave some discrepancy between the simple nanodisc model and the SAXS data. The structural models from SAXS and EM have the same overall dimensions, except for a larger ellipticity and a smaller circumference in the SAXS model compared to the EM model.

The SAXS model has an ellipticity of 1.5 ± 0.2, whereas the EM model has an ellipticity between 1.2 and 1.3. Polydispersity is known to affect a SAXS curve in a similar way as ellipticity. Polydispersity was observed, in the form of a mixed population of two-protein particles and a fraction of three-protein particles, as well as in variation in the number of lipids per nanodisc, and this polydispersity was not included in the SAXS model. Therefore, the ellipticity from the SAXS analysis may be slightly exaggerated to account for the lack of polydispersity in the model. This is not an issue in EM, as polydispersity and ellipticity are not correlated in the same way.

The circumferences calculated form the SAXS models were, respectively, 389 ± 30 Å for ApoE3-POPC and 355 ± 18 Å for ApoE3-DMPC (Table 5), and thus smaller than those obtained from EM, which were, respectively, 426 ± 22 Å and 413 ± 22 Å (Table 3). An ideal *α*-helix has a length of about 1.5 Å per residues, so ApoE3 (299 amino acids) was expected to span about 450 Å, which is close to circumferences of the EM models. The circumferences of the SAXS models are calculated from the semiaxes of the nanodisc and are therefore correlated to the ellipticity. The discrepancy in circumference might therefore also be caused by the fact that the geometrical SAXS model did not take polydispersity into account. The forward scattering of the model is still right despite the shorter protein belt. This may be because the protein belt is slightly higher (24 Å) than two stacked *α*-helices (2 x 10 Å).

### 3.2 ApoE3^80-255^ forms heterogeneous discoidal particles with four proteins per particle

EM data showed that the truncated construct ApoE3^80-255^ also forms disc-shaped particles, and the SAXS data can be fitted with a nanodisc model. Both EM, SAXS, native gel and SEC supported that the ApoE3^80-255^-based nanodiscs were larger than ApoE3-based nanodiscs. Phosphate analysis and SAXS held together supported a model with 4 proteins and about 300 lipids per disc, which was also consistent with the dimensions from EM. However, cross-linking did not reveal a band at 4 x ApoE3^80-255^. We suggest that 4-protein nanodiscs are formed, but that the proteins are organized pairwise around the central lipid bilayer, such that the relatively short crosslinking molecule does not connect the two pairs of the 4-protein nanodiscs effectively.

The ellipticity for the ApoE3^80-255^-lipid EM model was between 1.2 and 1.3 (Table 2), whereas the SAXS model had an ellipticity between 1.8 and 2.0 (Table 5). The same argument as for the ellipticity of ApoE3-lipid particles applies, and as the heterogeneity is larger for ApoE3^80-255^-lipid particles, the ellipticity of the SAXS model is also expected to be more severely exaggerated. Therefore, we expect the actual ellipticity to be closer to that of the EM envelope.

The circumferences of the SAXS models were about 450 Å (Table 5), and thereby smaller than the circumference of the EM models, which were about 550 Å (Table 2). Like lipid-bound ApoE3, the EM models were closest to the expected value. With two consecutive ApoE3^80-255^ proteins (2 × 199 residues) and 1.5 Å per residue for an ideal *α*-helix, the expected circumference was about 600 Å. Again, this discrepancy might be ascribed to the simplicity of the SAXS model, which, firstly, does not take polydispersity into account, and secondly, uses a ring with a rectangular cross section (10 Å x 24 Å) to model two stacked *α*-helices.

### 3.3 Dependency on lipid type

Our study demonstrates that the overall architecture of the particle does not change if the lipid changes from DMPC, which was in the gel phase in the present study, to POPC, which was in the fluid phase. Hence, we claim that lipid-bound ApoE3 forms rather stable disc-shaped HDL particles. Low density lipoproteins based on ApoE3, containing much larger amounts of lipids, cholesterol or triglycerides, may however have completely different morphologies.

### 3.4 Comparison with ApoE4-based particles

ApoE4 HDL particles have previously been described as ellipsoids with a twisted bilayer and a helical hairpin conformation of the ApoE4 [20]. We investigated protein-lipid particles formed with the more common isomorph ApoE3 (78 % allele frequency). Our model is also elliptical, and it has the same overall size as the ellipsoid model, but the bilayer is flat, resulting in a disc-shaped particle, rather than a 3-dimensional ellipsoid. We acknowledge that this structural difference may be an effect of the subtle difference between ApoE3 (Cys112 and Arg158) and ApoE4 (Arg112 and Arg158). The model for ApoE4-based lipid particles was also supported by SAXS, but we note that the authors ignored the scattering contribution from the lipid bilayer in their underlying SAXS data analysis. This is not a valid assumption, as the excess scattering length is non-zero for both lipid heads and lipid tails (Table 1), and indeed models with and without lipids leads to very different scattering curves as can easily be shown (Figure SI-2). This severely hampers the SAXS analysis leading to the ellipsoid model proposed by Peters-Libeu *et al*. [20]. In addition to this criticism, reported high values for the bending moduli of lipid bilayers also work against models with twisted bilayers [52]. This is further supported by a paper discussing a similar twisted model for ApoA1-lipid particles [53]. Here, two different MD approaches show the strong driving force towards a flat lipid bilayer. Hence, we find it more likely that lipid-bound ApoE4 also forms flat disc-shaped particles like those proposed here for lipid-bound ApoE3.

The central eight helical domains of ApoE3, i.e. the ApoE3^80-255^, were also found to self-assemble into soluble particles when associated with lipids. The particles are also discoidal, but slightly larger and more heterogeneous than the full-length ApoE3-lipid particles. Moreover, there were 4 copies of ApoE3^80-255^ per particle, instead of only 2 copies of ApoE3 in the ApoE3-lipid particles.

### 3.5 Structure of other apolipoproteins and the role of helical repeats

Inspired by the behavior of ApoA1, where ApoA1 with and without its globular domain form particles of basically the same size and shape, we hypothesized that the central ApoE3^80-255^ construct was the main domain for forming discoidal particles for ApoE3. As stated above, this turned out not to be the case for ApoE3 despite their similarities. The cause of this might be that the helical repeats in ApoE3 are only in one case punctuated by a proline residue, whereas seven out of ten helices are punctuated by prolines in ApoA1. This likely causes the central helices in ApoE3 to be more rigid and straight than those of ApoA1, requiring the end domains of ApoE3 to fine-tune the self-assembled particles. Our data and models however support the notion that the amphipathic helical repeats found in ApoA1 and ApoE3 might be a common denominator for the ability to solubilize lipids and for forming flat discoidal particles.

A flat overall discoidal morphology has also been observed for HDL particles containing ApoA1, ApoA2 [54], ApoA4 [55], and ApoA5 [55]. Going beyond the full-length apolipoproteins, similar structures have been reported for the 22 kDa N-terminal fragment for ApoE3-DMPC [56] and even helical repeat mimicking peptides [50,51,57]. Accordingly, it could be beneficial to revisit the structure of some of these lipoproteins (e.g. with ApoA2, ApoA4, and ApoA5) and investigate their self-assembly and formed structures based on duplicated domains and exon structure from genomic data.

### 3.6 Conclusion

Through a combination of SEC, native PAGE, crosslinking, phosphate analysis, negative stain EM and SAXS we find that full-length ApoE3 as well as truncated ApoE3 (ApoE3^80-255^) self-assemble with both DMPC and POPC to form disc-shaped HDL particles, or so-called nanodiscs. The particles with ApoE3^80-255^ are largest, with about 300 lipids and with the major population having four proteins per particle, whereas particles with full-length ApoE3 contain about 200 lipids and has two proteins per particle.

## 4 Acknowledgements

We thank for the beamtime allocated the EMBL P12 SAXS beamline and thank Haydyn Mertens for support. We thank for electron microscope time at the Core Facility for Integrated Microscopy (CFIM) at University of Copenhagen and at EMlab at Aarhus University. We thank Thomas Boesen, Nanda Gowtham Aduri, Klaus Qvortrup and Tillmann Pape for invaluable contributions with obtaining and processing the EM data. Finally, the authors acknowledge co-funding from the Lundbeck Foundation Brainstruc project and a Lundbeck Foundation International Postdoc grant to SRM, the CoNeXT programme and the Carlsberg Foundation, the Novo Nordisk Foundation Synergy project, and the KU2016 BioSynergy programme.

## 5 Author Contribution

Conceptualization: AHL, LA, SRM; sample preparation: AHL, NTJ, SRM; SAXS data collection: AHL, SRM, NTJ; EM data collection: AHL, NTJ, SRM; SAXS analysis: AHL, EM analysis: AHL, MG; original draft: AHL, SRM; review and editing: AHL, NTJ, LA, SRM.

## 7 Supplemental Information

### 7.1 List of supplemental tables and figures

Table SI-1. Protein sequences.

Figure SI-1. Guinier plots of the SAXS data.

Figure SI-2. Comparison of SAXS models with and without lipids taken into account.

### 7.2 Table SI-1. Protein sequences

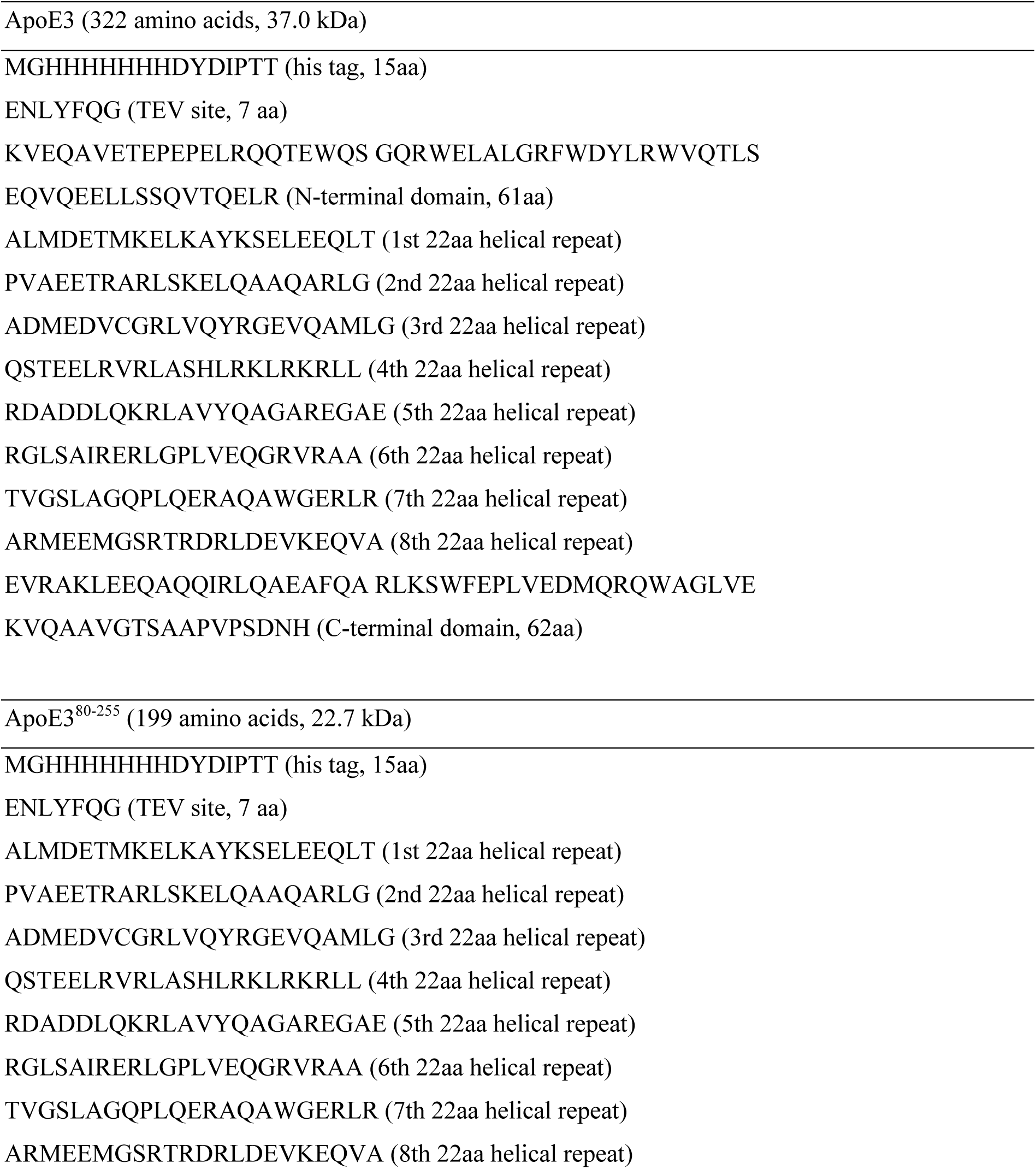

### 7.3 Figure SI-1. Guinier plots of the SAXS data

All SAXS data decays linearly on a Guinier plot (logarithm of the intensity against the square of the momentum transfer). That behavior is a good indication that there are no aggregated particles in the sample.

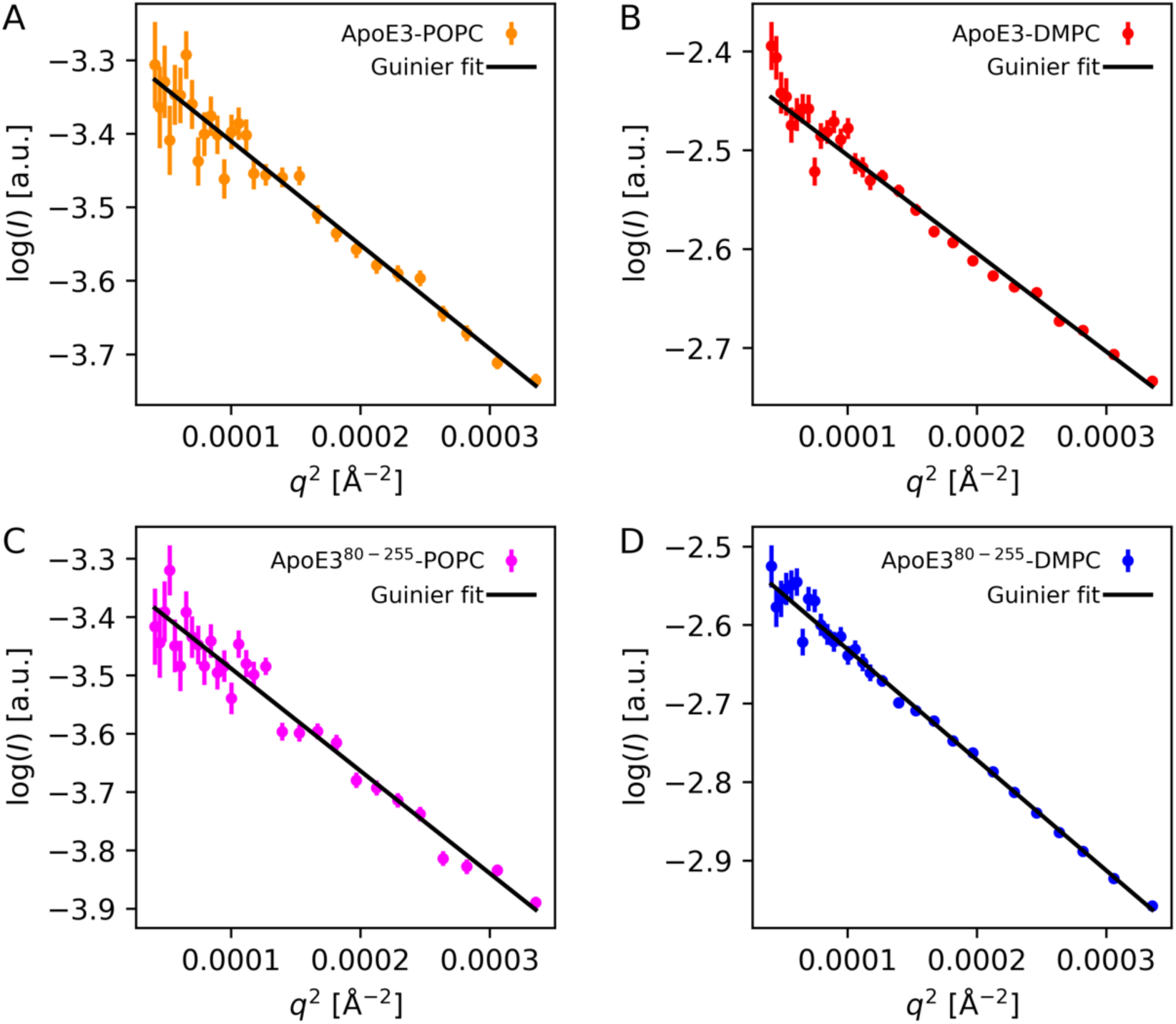

### 7.4 Figure SI-2. Comparison of SAXS models with and without lipids

SAXS data of ApoE3-DMPC particles (red) fitted with a model of nanodiscs, where the DMPC lipids have been included (black), and a model with the same parameters, but ignoring lipids (grey). DMPC has an average electron density very close the solvent (Table 1), but as the electron density of the lipid headgroups and tail groups differ from the solvent density (Table 1), the scattering contribution from DMPC cannot be ignored.

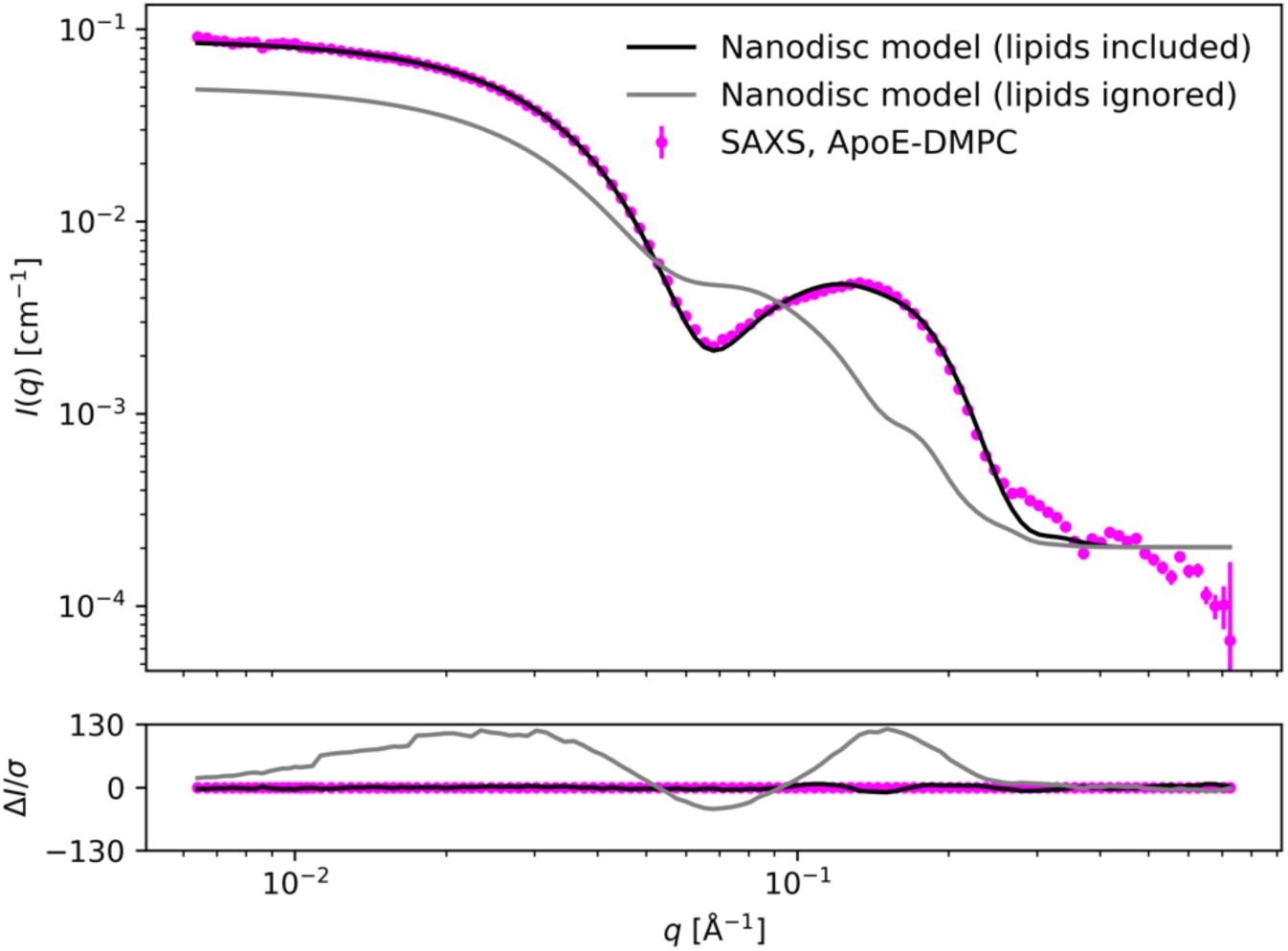

